# Mathematical modeling of the protective coloration of animals with usage of parameters of diversity and evenness

**DOI:** 10.1101/822999

**Authors:** Yu. Bespalov, K. Nosov, O. Levchenko, O. Grigoriev, I. Hnoievyi, P. Kabalyants

## Abstract

Adaptive mechanisms performing at different levels of organization of living matter play an important role in theoretical biology. One of the important cases of such mechanisms is the protective coloration of animals, that masks them on the ground.

The article aims at building mathematical models of the performance of the protective coloration of animals, depending on the specific situations of their adaptation to a particular area. The results of the study can be used to create remote technologies for detecting animals of certain species at a considerable distance.

## Introduction

The study of adaptive mechanisms at different levels of organization of living matter is one of the most important topics of theoretical biology. At the same time, the results of these studies in many cases have important relevance. To the full extent, this relates to one of the manifestations of adaptive mechanisms—protective coloration of animals.

Global climate changes cause the urgency of both these aspects of biological science. It should be noted that global climate changes, in general, pose many severe problems of the existence of human civilization. Among the problems related to biology are problems arising in the agricultural sector of the world economy. In particular, they make urgent the problems of creating and using new technologies of cattle breeding. We bear in mind the technologies efficient in large areas, the climate of which will become warmer and arid, but the development of the infrastructure on them for the life of a significant permanent population is unpractical. In such situations, the efficiency of the agricultural sector of the economy can be implemented through the development of cattle breeding with use of high-tech methods, for use of which a small in number staff is sufficient, for remote monitoring the state of numerous herds of animals in vast, sometimes difficult-to-reach areas. Such methods use aviation, as in the case of Australia (Mulero-Pázmány et al., 2017), including drones. We are talking about the usage of drones for controlling the movement and condition of grazing animals as well as predators and competitors, which negatively affect the grazing animals’ productivity. Organisms—indicators of the ecological state of grazing places—may also be controlled in such a manner. For example, the disappearance of shes or other animals in pounds and streams used for stock watering can serve as indicators of an poor state of their ecology.

Effective application of aviation may be prevented by the protective coloration of domestic and wild animals. For example, Australian camels belong to both of these categories.

In particular, this problem will arise during the wide economic use of relatively recently domesticated animals, which may prove economically advantageous in the described situations associated with global warming. Common elands (*Taurotragus oryx*) in Askania-Nova National Park of Ukraine can be mentioned as an example. The protective mechanisms of animal coloration are very complex and are systemic in nature. This implies the use of certain approaches to mathematical modeling procedures for their study (Turing, 1952; Murray, 1981a,b; Murray and Maini, 1986; Fennell et al., 2019). This paper aims to study the abilities of usage of new approaches described below.

## 1 Literature Analysis and Statement of the Problem

For a long time, the coloration of animals has been of interest to representatives of different branches of science (Turing, 1952; Murray, 1981a,b; Murray and Maini, 1986; Endler and Mappes, 2017; Fennell et al., 2019). They investigate multiple aspects of this problem and specified objects of the study.

The impact of certain optical phenomena inherent in the underwater environment on masking adaptation mechanisms of pelagic shes is studied in (Johnsen et al., 2014).

The mechanisms that marine animals use for masking bioluminescence and polarization of light are described in (Marshall and Johnsen, 2017; Feller et al., 2017).

The work (Lind et al., 2017) is devoted to the laws of the co-evolution of color vision and animals’ coloration.

In (Shawkey and D’Alba, 2017), the relationships between the mechanisms of production of substances determining the coloration of animal substances and their use were analyzed.

The parameters of the genome of their population affecting the coloration of wild animals were analyzed in (San-Jose and Roulin, 2017).

The mechanisms of performance of the silhouette of animals’ coloration (camouflage) that divide the visual perception, as well as the conditions of its adaptability and environmental significance are investigated in (Merilaita et al., 2017; Duarte et al., 2017).

In particular, the role of coloration of boundary areas of the silhouette for the destruction of a holistic visual perception of the contour of animals is described in (Merilaita et al., 2017).

The work (Nothdurft, 2018) is devoted to the processing of visual information by the brain. We are talking about aspects that affect the speed of various processes of visual perception.

The role of diversity at different levels of organization of living matter in the performance of protective animals’ coloration is an important problem.

In this regard, it should be noted that the problem of the role of diversity in the performance of living systems is one of the essential problems of fundamental biology. The results of the study of this problem by environmentalists during many decades are presented in the works (Bukvareva and Aleshchenko, 2012, 2013, 2005), devoted to the principle of optimal diversity and research, in particular, by methods of mathematical modeling, manifestations of this principle.

The protective coloration of animals is an important and, moreover, relatively convenient for study, case of the performance of adaptive mechanisms. In this case, the relationship between diversity of the system and the character of its performance plays an important role. (At the organismic, but not only at the organismic, level of organization of living matter.) Mathematical modeling as an effective tool for these mechanisms’ study.

In this regard, works (Turing, 1952; Murray, 1981a,b; Murray and Maini, 1986) by A. Turing based on the model (Turing, 1952) play an important role.

More recently, specific mechanisms that implement such models at the molecular level of the organization of living matter were described (Kondo and Miura, 2010; Sheth et al., 2012). We bear in mind the implementation of parameters of the periodicity of not only color but also other aspects of morphology.

In the work (Feller et al., 2017), the authors present the results of using neural networks for the study of the performance of the protective coloration of animals. In this work, it is noted that in the case of usage of neural networks the requirements for the initial data are not so strict as for Big Data, but still quite serious.

A certain statement confirmed in (Endler and Mappes, 2017; Johnsen et al., 2014; Marshall and Johnsen, 2017; Feller et al., 2017; Lind et al., 2017; Shawkey and D’Alba, 2017; San-Jose and Roulin, 2017; Merilaita et al., 2017; Duarte et al., 2017; Nothdurft, 2018) appears to be well-grounded. It means the following. In cases when the coloration of animals serves to protection and masking, its formalized description requires approaches based on the Turing’ model (Turing, 1952; Murray, 1981a,b; Murray and Maini, 1986) supplemented by others ones.

Approaches that enable to simulate the aspects of protective coloration of animals providing the adaptation to the diversity of colorimetric parameters (CPs) of their habitats should play an important role. In particular, we are keep in mind the adaptation to the diversity of plant communities in these places. Global climate changes will frequently enough create situations requiring the adaptation of animals’ coloration to a high level of diversity of these CPs.

On the other hand, there might be situations creating threats to biosafety, which require decision-making in case of an acute lack of time and resources. This could include a lack of time and resources for collecting initial data for performance of decision support systems (DSS). So, mathematical approaches allowing one to develop DSSs that work with relatively small volume of data arrays are topical. This also relates, in particular, to remote methods for obtaining information about the localization and migrations of animals having masking coloration. So, the development of sufficiently universal unmasking technologies will be required. It may be, in particular, technologies based on certain types of mathematical models of mechanisms of adaption of protective coloration of animals to diversity of their habitats. To be exact, we talk on types of mathematical models that do not require a large amount of initial data.

In (Bespalov et al., 2017; Balym et al., 2017; Vysotska et al., 2017; Bespalov et al., 2018; Nosov et al., 2018), a new approach that satisfies this requirement to mathematical modeling of the protective coloration of animals is proposed. It enables to use the data with gaps and having a minimum volume. (We are talking on the number of observations that makes it possible to correctly calculate the correlation matrix between the components of a certain system). Then, with use of Discrete Models of Dynamic Systems (DMDS), on the basis of such a matrix, the structure of inter-component and within component relationships in the system can be built, due to the positive and negative effects of the components on each other and on themselves (Zholtkevych et al., 2013). On the base of this relationships structure, an idealized trajectory of systems (ITS) can be calculated (Bespalov et al., 2017; Balym et al., 2017; Vysotska et al., 2017; Bespalov et al., 2018; Nosov et al., 2018). This trajectory represents the dynamics of the system. In this particular case, the ITS can represent a cycle of values’ change of colorimetric parameters of plant communities in animal habitats. An idealized pseudo-trajectory of the system (IPTS) can also be built. It reflects a set of CPs values of the disruptive (camouflage) protective coloration of an animal that provides adaptation to a specific area. (But this trajectory do not reflect the real time dynamics.) In (Bespalov et al., 2017; Balym et al., 2017; Vysotska et al., 2017; Bespalov et al., 2018; Nosov et al., 2018), it was shown that animals possessing such a protective coloration can be unmasked using procedures that require only initial data for their implementation, which can be obtained by computer analysis of the components of the RGB model of digital imaginary. (In a number of cases, we can talk about digital imaginary obtained with the help of the equipment included in the package of delivery of relatively cheap and easy-to-use drones.) These methods are based on a comparative analysis of ITSs and IPTSs that was built with use of DMDS. This comparative analysis results in revealing the differences in diversity of values of CPs of plant background and coloration of animals. The differences are used later for unmasking. We note once again that the protective coloration of animals is a special case of the performance of adaptive mechanisms. In this case, the role of the diversity of certain manifestations of this adaptive mechanism with the nature of its performance was investigated. In (Bespalov et al., 2018), using antelopes *Taurotragus oryx* as a case study, it was shown that, along with this aspect of the diversity of the living system, in this case, the degree of evenness of the values of CPs of protective color also plays a role. It should be noted that a similar systemic effect has been proposed a long time ago for use by ecologists in the analysis of the structure of biological communities (Margalef, 1968).

We are talking on the analysis based on the Shannon index (Shannon, 1948). It seems appropriate to analyze the aspects of diversity and evenness of the protective coloration of animals, taking into account the specific circumstances of their adaptation to a particular area.

The aim of the current study is to build mathematical models of performance of the protective coloration of an animal that depend on the specific circumstances of their adaptation to a particular area. Specifically, we build the mathematical models using the parameters of the diversity of CPs’ values and their evenness.

In the course of such modeling, the possibility of applying the obtained results for remote registration of animals in areas of certain types was taken into account.

To achieve this aim, the following tasks were solved:

- to build the models of the animals’ protective coloration and the spatial-temporal distribution of CPs of local vegetation in the form of scatter plots in feature spaces with coordinates that reflect the diversity and evenness of the values of the corresponding CPs;
- to develop the image processing procedures for diagnostics of the presence or absence of certain animals in certain types and sections of the terrain, on the basis of these models.

## 2 Obtained Results

This work results in the models of the protective coloration of animals and CPs of plant communities in their habitats. These models have the form of scatter plots in a 2D feature space (2DFS). Coordinates of this 2DFS were selected in accordance with results of modeling presented in (Bespalov et al., 2018), with use of DMDS, systemic parameters of performance of the protective coloration of animals. We are talking about the system colorimetric parameters (SCPs) and primary colorimetric parameters (PCPs) obtained by computer analysis of components of the RGB model of digital imaginary of animals on the background of vegetation of their habitats. To find the values of SPCs and PCPs, relatively large fragments of images (hereinafter referred to as macrosegments) were divided into segments. The segments, in turn, were divided into microsegments. We analyzed SCPs of segments containing full or partial images of animals and the segments containing only background images of plant communities in animal habitats.

The values of following CPs reflecting the tones of animals’ coloration and its evenness were calculated for each microsegment:

- the value of 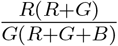;
- the value of 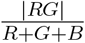.

For a set of microsegments of any segments, the following values of SCPs were calculated:

- the range of 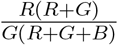;
- the range of 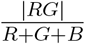;
- the mean of 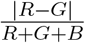.

These values reflect the average values of evenness uniformity and diversity of tones of animals’ coloration.

These system colorimetric parameters were used as coordinates of the 2DFS in which these scatter plot were built. These parameters reflect the variety of combinations of CPs’ values and their evenness.

Scatter plots of parameters of protective coloration of feral one-humped camels (*Camelus dromedarius*), dingo dogs (*Canis lupus dingo*), and kangaroos (*Macropus rufus*), as well as plant communities in the regions of Australia where these animals live, were built in such two-dimensional space In addition, in the same feature space the scatter plots of protective coloration of shes (*Cyprinus carpio*), as well as plant communities of ponds in Ukraine, where they are growing, were built. These scatter plots are shown in Figs 1, 2, 3, 4, 5, 6, 7, 8.

**Fig. 1:**
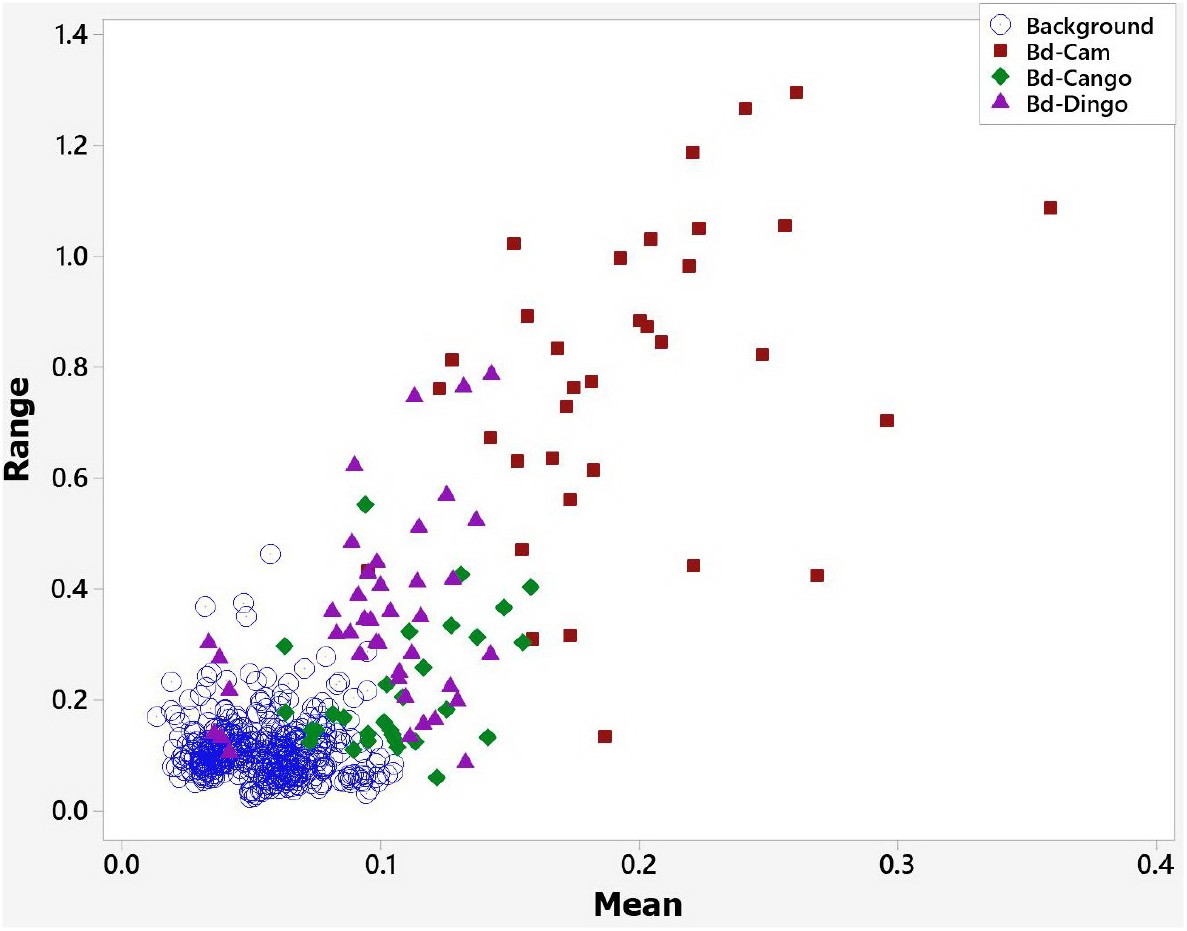
Scatter plot of SCPs of segments with bodies of different animals and background. Mean: the mean of 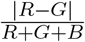; Range: the range of 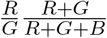; Bd-Cam: body of camel; Bd-Cango: body of kangaroo; Bd-Dingo: body of dingo; Background: background without animals

**Fig. 2:**
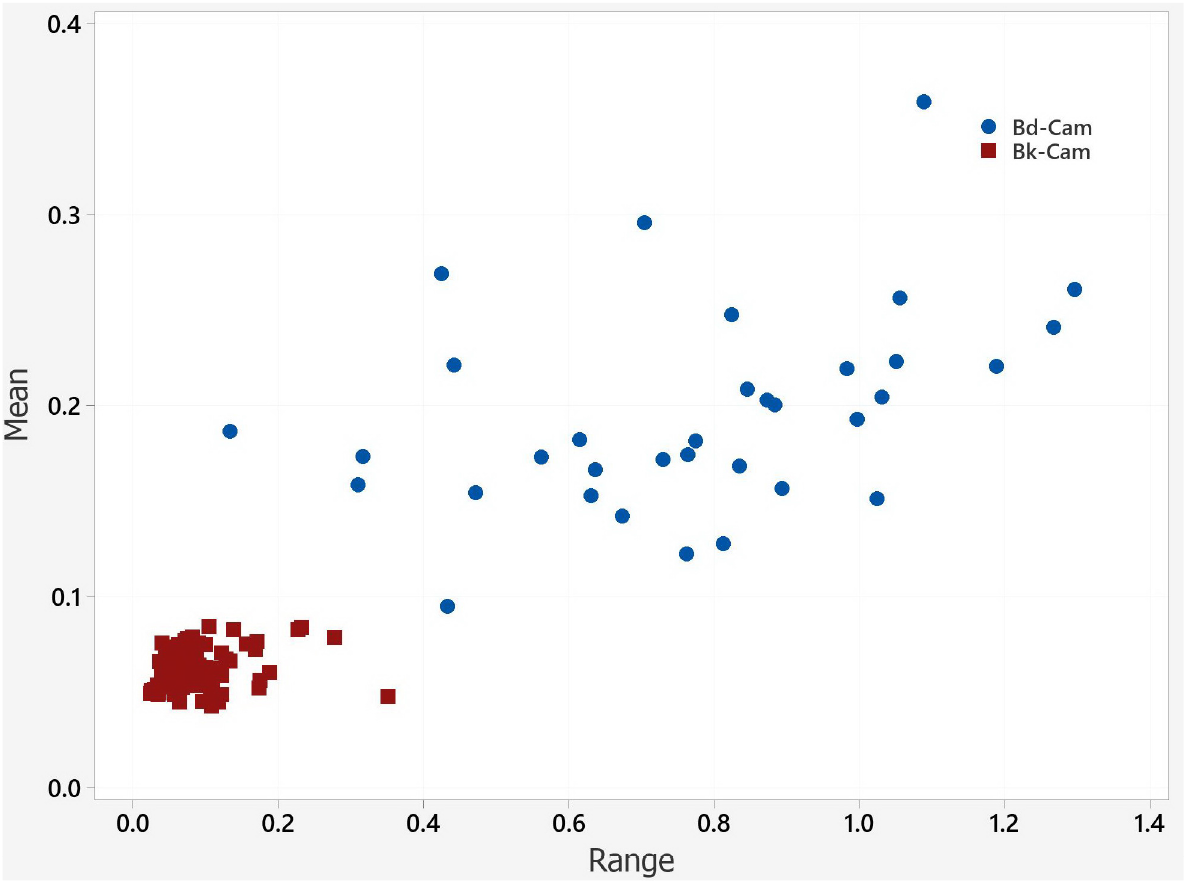
Scatter plot of SCPs of segments for camels. Range: the range of 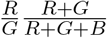; Mean: the mean of 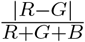; Bd-Cam: body of camel; Bk-Cam: background without animals

**Fig. 3:**
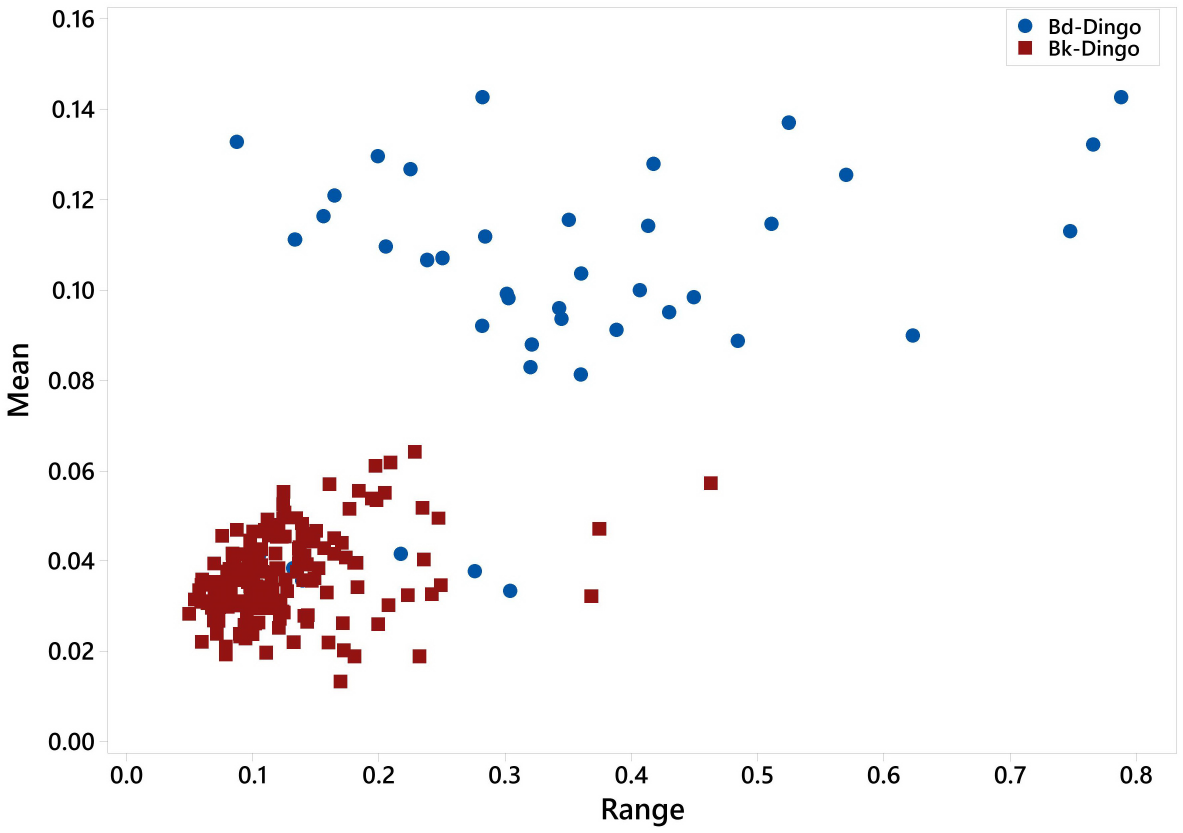
Scatter plot of SCPs segments for dingos. Range: the range of 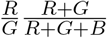; Mean: the mean of 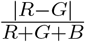; Bd-Dingo: body of dingo; Bk-Dingo: background of dingo

**Fig. 4:**
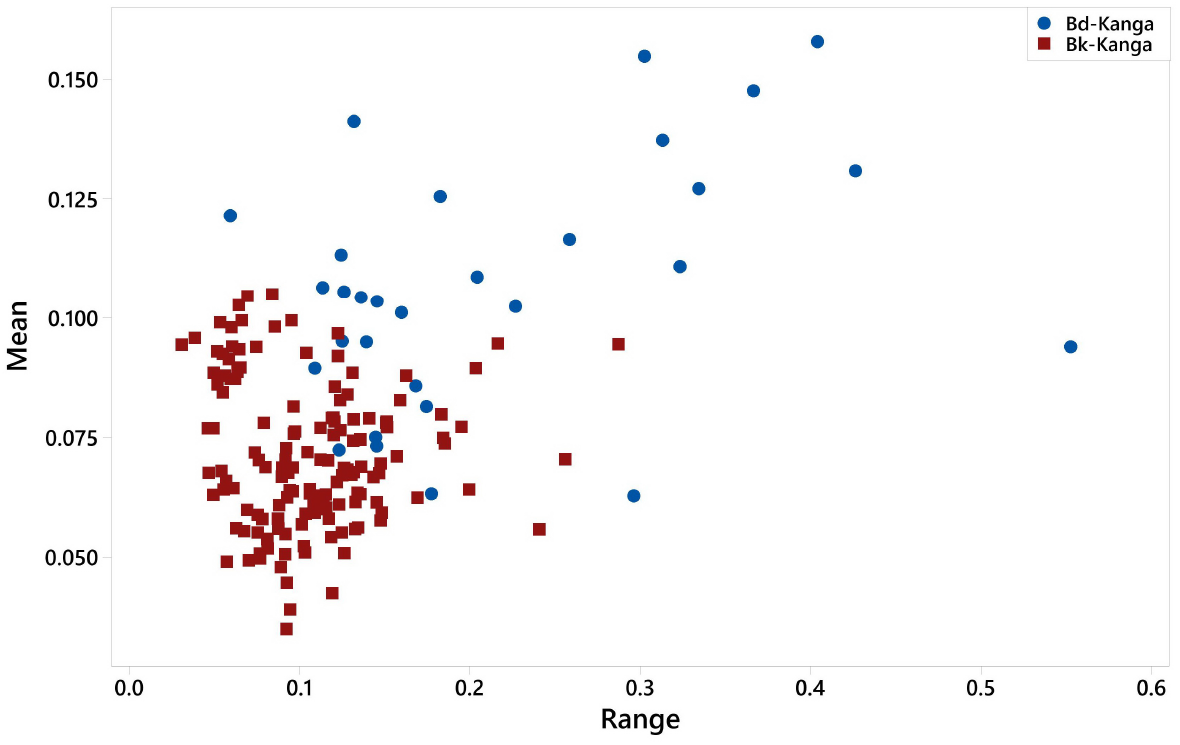
Scatter plot of segments for camels. Range: the range of 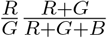; Mean: the mean of 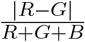 Bd-Kanga: body of kangaroo; Bk-Kanga: background of kangaroo

**Fig. 5:**
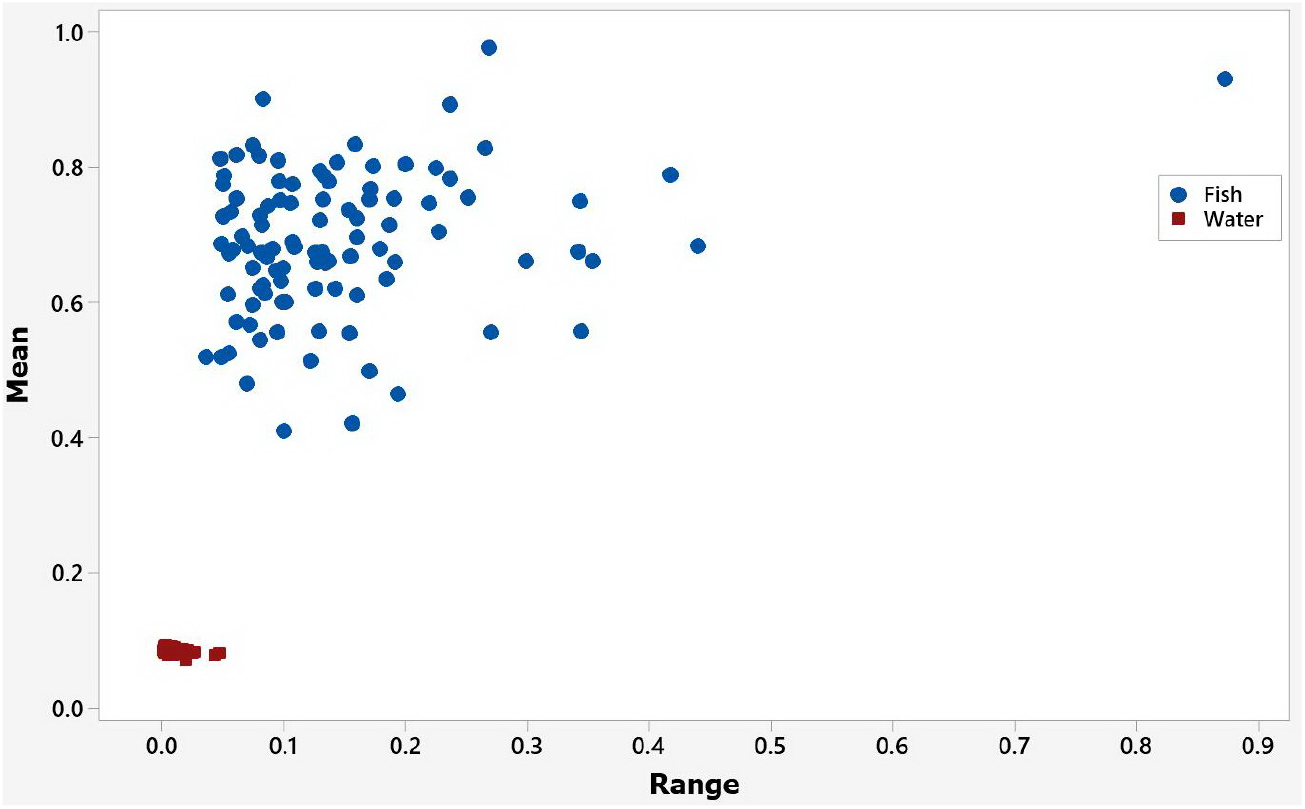
Scatter plot of segments for carps. Range: the range of 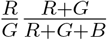; Mean: the mean of 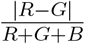; Fish: segments with bodies of karps; Water: segments without karps

**Fig. 6:**
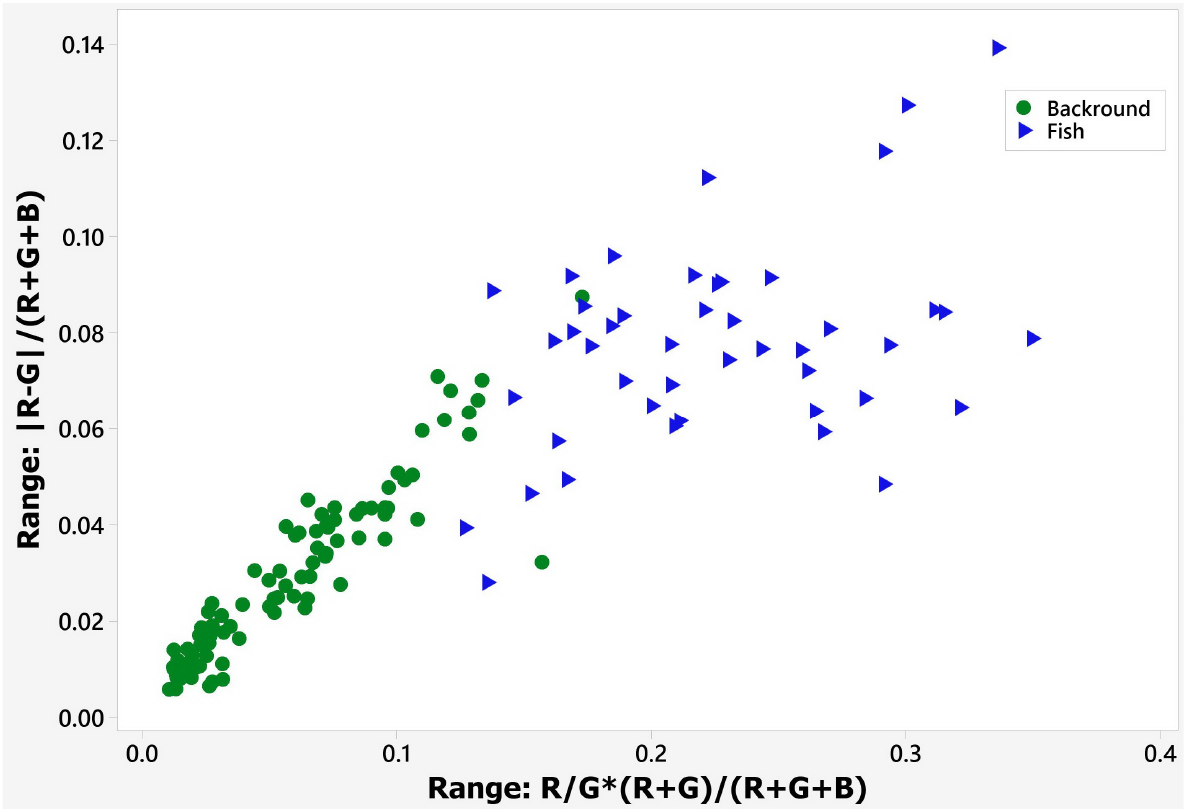
Scatter plot of segments for carps. Axis *X*: the range of 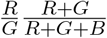; axis *Y*: the range of 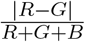; Fish: segments with bodies of carps; Background: segments without carps

**Fig. 7:**
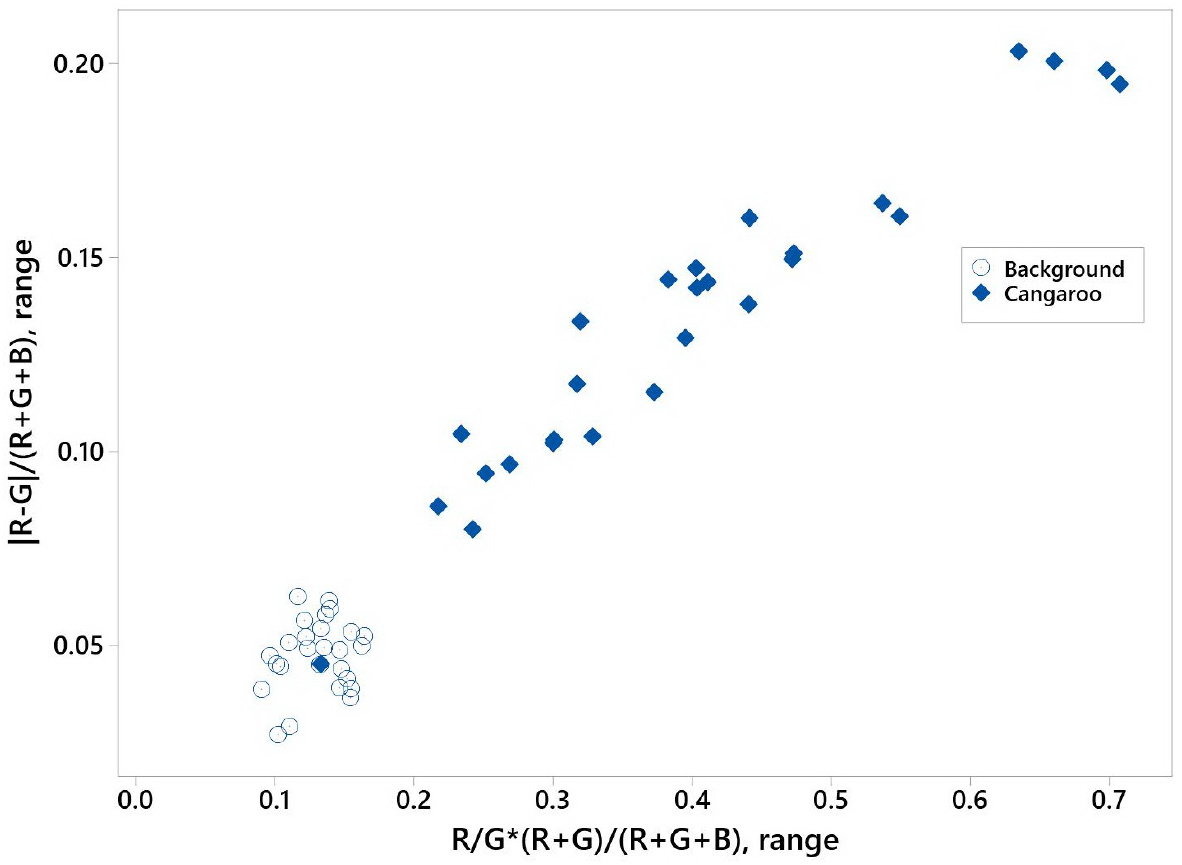
Scatter plot of segments for kangaroos. Axis *X*: the range of 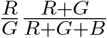; axis *Y*: the range of 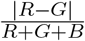; Cangaroo: segments with animals; Background: segments without animals

**Fig. 8:**
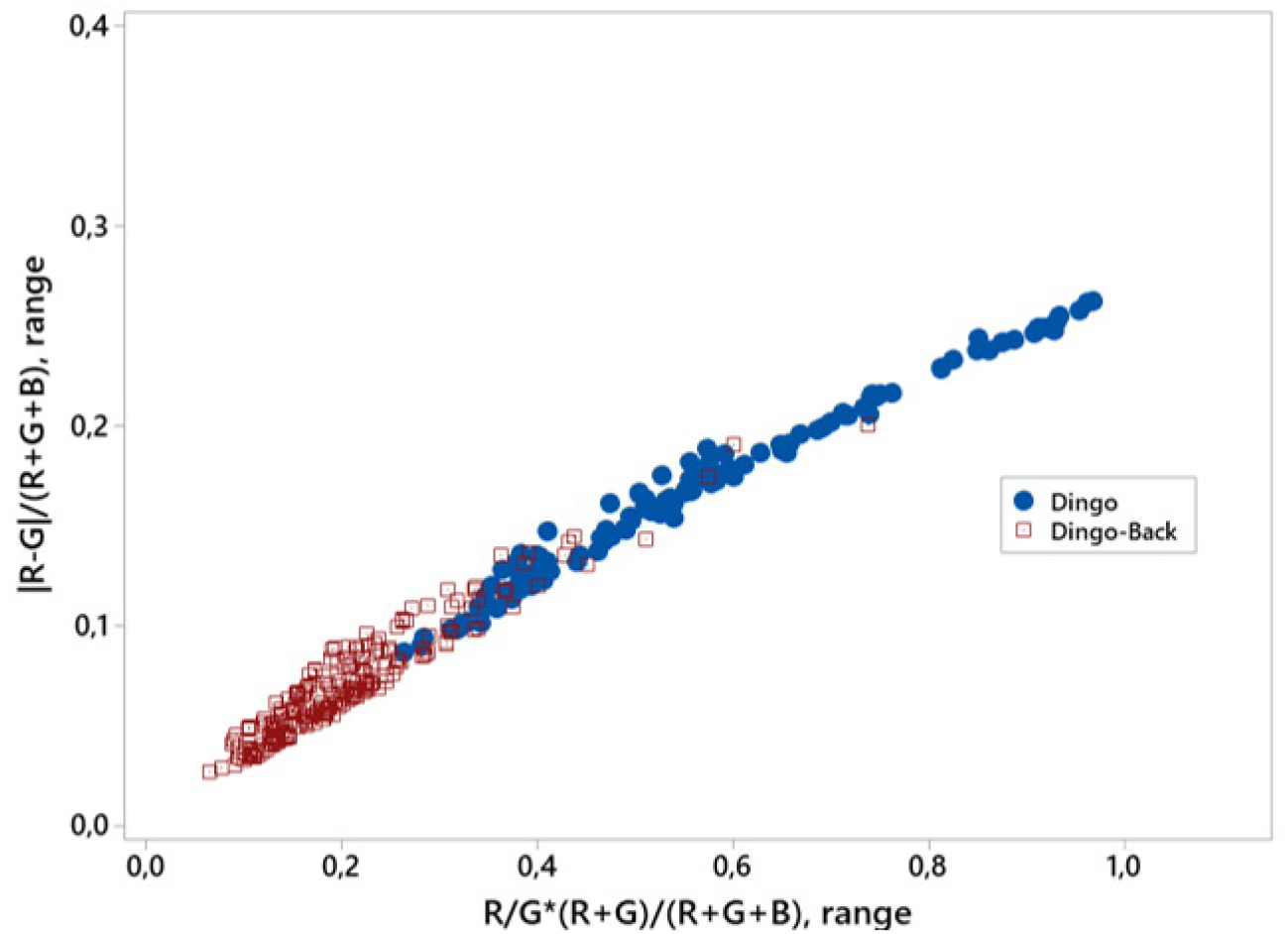
Scatter plot of segments for dingos. Axis *X*: the range of 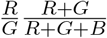; axis *Y* axes and the range of 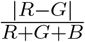; Dingo: segments with animals; Dingo-Back: segments without animals

A comparative analysis of the scatter plots presented in Fig. 1-4 and 5-8, gives some reasons for formulating a rule to which the parameters of diversity and coloration evenness follows. These parameters depend on the situation (evolutionary history) in which this adaptive trait was formed. More specifically, these measures consist of: The period of time from the appearance in Australia of camels (*Camelus dromedarius*), dingo dog (*Canis lupus dingo*) and kangaroo (*Macropus rufus*) may be arranged in an ascending order. The values of above-mentioned system colorimetric parameters used in scatter plots can be also ranged in decreasing order. Namely:

- the range of the value 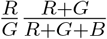;
- the range of the 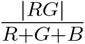;
- the mean of 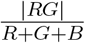.

This effect can be explained in such a way.

Camels (*Camelus dromedarius*) appeared in Australia about two centuries ago. Their wild or, rather, feral form began to spread in desert and semidesert regions of the continent almost at the same time. Over the specified period of time, the Australian population of this species has greatly increased its number and range. The ever-growing range includes various (albeit very similar) plant communities. The colorimetric parameters of these communities are distinguished by a significant variety of tones. The ongoing and currently expanding range suggests the possibility of increase of this diversity. It is mainly about tones of green and red, reflecting the ratio of green chlorophyll to red-yellow-orange plant pigments. This ratio is significantly different for different plant communities and different stages of their development. In such a situation, the maximum possible variety of its shades will meet the optimal strategy for performance of animals’ protective coloration. We mean the adaptive function of color enabling to destroy the integral visual perception of the animal’s silhouette. This suggests a certain law of diversity of tones of silhouette’s fragments. The essence of the law is as follows. At any point of time and space, there is a number of silhouette’s fragments merging by color with the plant background sufficient for destruction of silhouette’s perception. (Or, at least, the diversity of tones of these fragments should be commensurately to the diversity of CPs of the plant background at specified points in space and time.) In the case under investigation, the measure of this diversity is the range of the value 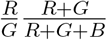 calculated on the set of microsegments of each segment. (The microsegments correspond to the above mentioned fragments of the animal’s silhouette. For animals having the size approximately equal to human’s one were about several centimeters, by analogy with camouflage elements of army uniforms or spots of the coloration of leopard, jaguar, cheetah.)

It should be noticed that an increase in the diversity of tones destroying the integral perception of the silhouette is related to the consumption a certain resource. More specifically, it means the following. An increase in the number of tones implies a decrease in the angular size of spots of different colors on the body of an animal. This decrease can lead to merging the visual perception of these spots. Such merging will lead to elimination of the effect of destruction of integral perception of the silhouette.

When this resource is limited (e.g., upon decrease of the animal’s size and increasing the distance of its visual perception), the evenness of the protective coloration gets adaptive meaning that increases its universality. The brown coloration of many animals may be referred as an example. It provides the merging of silhouette’s fragments against a plant background accompanying by relative predominance of both green chlorophylls and red-yellow-orange pigments. In the case under study, high evenness of CPs corresponds to the low value of 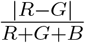.

Not only the evenness matters, but also its range and diversity are important. In the case under study, the measure of this aspect of diversity is the range of 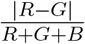. To a certain extent, the diversity of evenness values of CPs can be used as a measure of permissible dissimilarity of colors of the animal’s coloration and plant background. (The degree of this permissible dissimilarity is determined by the development of color vision in animals belonging to the same trophic network or chain, lighting conditions and other factors. In the current study, we do not consider these conditions.)

Dingos staying in Australia for thousands of years occupy an intermediate position between camels and kangaroos according to the range of 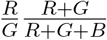. Kangaroos formed as a species in Australia and adapted, due to long-term natural selection, to ecological niches of this continent have the lowest values of this indicator. The evenness of the CP of dingos and kangaroos is approximately the same and significantly higher the camels’ CP. The range of the evenness of kangaroos and dingos is also quite close and significantly lower than that this indicator of camels.

This corresponds to the strategy for performance of protective coloration of animals, which have for a long time occupied certain ecological niches, but which adaptive mechanisms work in conditions of changing ecological niches. This is possible due to the relative universality of these adaptive mechanisms.

The scatter plots in Fig. 5, 6 built for carps (*Cyprinus carpio*) living in Ukrainian sh ponds reflects a strategy of coloration of animals that have found an ecological niche for themselves for a long time ago. But, later, this coloration, as well as the ecological niche, has significantly changed under influence of human activity. Moreover, these changes reduce the role of coloration in the adaptation of animals to the environment.

The differences in the values of SCPs of sh and plant communities of water of the ponds presented in Fig. 5, 6 are important for considering the practical results of this work. The differences in the values of SCPs of camels, dingoes and kangaroos and plant communities in their habitats presented in Fig. 1-4 and Fig. 7-8 are also important. The differences in SCPs of protective coloration of different Australian animals shown in Fig. 1-4 and Fig. 7–8 have also certain importance.

In accordance with the statement in (Nosov et al., 2018), the diversity of protective coloration of an animal with a rather small probability will be larger in comparison to whole color diversity of its environment. So the probability of fact that the diversity of the protective coloration of animals will be greater than the diversity of its environment at a specify point of space and time is sufficiently large. Accordingly, SCPs, which are a measure of the color diversity of an animal, can be used to unmask it. In accordance with the results presented above here and, earlier, in (Bespalov et al., 2018), SPCs serving a measure of coloration evenness can also be used to unmask an animal. Having kept this in mind, we have advanced a working hypothesis according to which the usage of classification procedure on the base of neural networks enables unmasking the animals. We used classification procedures with the following colorimetric parameters as independent parameters:

- the range of 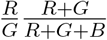;
- the range of 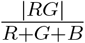;
- the mean of 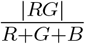.

These values reflect the average values of evenness uniformity and diversity of tones of animals’ coloration.

Testing this working hypothesis on the image of carps and Australian animals gives the following results. In Table 1 the results of animals/background recognition are presented. We used classification procedures on the base of neural networks for building the classifier, which distinguishes the segments with animals’ bodies and background. In Table 1 the following noti cations were used: **SNBd**: the **S**egments’ Number with animals’ **B**o**d**ies; **EBd**: number of **E**rroneously classified segments with bodies (including percentage); **SNBk**: the **S**egments’ **N**umber with **B**ac**k**ground; Erroneously classified segments with **B**ac**k**ground (including percentage).

**Table 1:** Results of recognition of segments with animals and background

| Species | SNBd | EBd (%) | SNBk | EBk (%) |
| --- | --- | --- | --- | --- |
| Carps | 428 | 11 (2.6) | 35 | 2 (5.7) |
| Kangaroos | 308 | 20 (6.5) | 55 | 0 (0.0) |
| Dingo | 53 | 1 (1.9) | 99 | 5 (5.1) |

## 3 Discussion

The results of this work are considered by the authors just as preliminary. They give, in our opinion, the basis for the following statements.

Certain parameters of diversity and evenness play a role in performing the adaptation mechanisms that disrupt the silhouette (camouflage) of animals’ coloration. In this regard, the results of (Bespalov et al., 2017, 2018) can be confirmed and obtain further development. The works (Bespalov et al., 2017, 2018) and the present paper deal with the diversity and evenness of tones of different fragments of the body’s coloration of an animal related to the nature of plant communities in their habitats. Namely: They deal with the diversity of combinations of the presence of red and green components and the evenness of their ratio, as well as the diversity of degrees of this evenness. The ratio of the red and green components in the coloration of an animal is related to the ratio of green chlorophylls and yellow-orange-red pigments in different areas and at different stages of development of these plant communities. Apparently, the possibility of extending the systemic effects described in this paper to other aspects of animal coloration, aspects of their morphology, and performance of adaptive mechanisms cannot be excluded.

In this work that includes a specified case study with Australian animals a certain regularity of the dependence of the role of diversity and coloration evenness of animals on the situation (evolutionary history) of their adaptation to ecological niches was described.

In this paper, this regularity performs as follows. The main role in the protective coloration of animals, which process of adaptation to new ecological niches is far from complete, plays the diversity of tones. In the paper’s case study, we investigated a wild (feral) Australian camel (*Camelus dromedarius*). When the process of adaptation to habitat in certain ecological niches continues for a long time (millennia, as in the case of *Canis lupus*), evenness parameters play an important role. Namely: in comparison with the rst case, against the background of a decrease in the variety of combinations of values and the brightness of red and green and the diversity of values of the evenness parameter 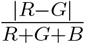), we see an increase in the average value of the parameter.

For kangaroos (*Macropus rufus*), a biological species that have arisen and evolved in Australia, these trends are even sharper.

The described regularity is illustrated by examples of animals, adaptation of which to ecological niches are extremely and obviously different in time. It is doubtful that the nature of this regularity is universal or, at least, significantly general. Nevertheless, we hope that the models describing this regularity (in a form of scatter plots) can be useful in some cases. For example, such models can help in using the parameters of diversity and coloration evenness of animals for their detection on area.

The analysis of the models presented here in the form of scatter plots enables to advance working hypotheses regarding recognition procedures for areas where the presence of animals are highly likely. Accordingly, for animals’ detection, based on the results of applying these procedures, video-information about these areas of the area can be automatically selected for veri cation by other methods: including with the participation of zoologists.

Recognition procedures that use these working hypotheses can be developed with usage of a set of mathematical tools. In this paper, we used neural networks for such procedures.

We can deduce that the results obtained in this work are reasonably sufficient for their application and implementation in procedures for recognition of areas with possible animals’ presence.

## References

Balym Y, Vysotska O, Pecherska A, Bespalov Y (2017) Mathematical modeling of systemic colorometric parameters unmasking wild waterfowl. Eastern-European Journal of Enterprise Technologies 5(2 (89)):12–18

Bespalov Y, Nosov K, Kabalyants P (2017) Discrete dynamical model of mechanisms determining the relations of biodiversity and stability at different levels of organization of living matter. bioRxiv DOI https://doi.org/10.1101/161687

Bespalov YG, Nosov KV, Kabalyants PS (2018) A mathematical model of the effect of natural selection on adaptation forms that implemented by disruptive coloration of Taurotragus oryx. bioRxiv DOI http://dx.doi.org/10.1101/368084

Bukvareva EN, Aleshchenko GM (2005) A principle of optimal diversity of biological systems. Uspehi sovremennoi biologii 125(4):337–347

Bukvareva EN, Aleshchenko GM (2012) The principle of optimal biodiversity and ecosystem functioning. International Journal of Ecosystem 2(4):78–87

Bukvareva EN, Aleshchenko GM (2013) Optimization, Niche and Neutral Mechanisms in the Formation of Biodiversity. American Journal of Life Sciences 1(4):174–183

Duarte RC, Flores AAV, Stevens M (2017) Camouflage through colour change: mechanisms, adaptive value and ecological significance. Philosophical Transactions of the Royal Society of London Series B, Biological Sciences 372(1724)

Endler JA, Mappes J (2017) The current and future state of animal coloration research. Philosophical Transactions of the Royal Society of London Series B, Biological Sciences 372(1724)

Feller KD, Jordan TM, Wilby D, Roberts NW (2017) Selection of the intrinsic polarization properties of animal optical materials creates enhanced structural reflectivity and camouflage. Philosophical Transactions of the Royal Society of London Series B, Biological Sciences 372(1724)

Fennell JG, Talas L, Baddeley RJ, Cuthill IC, Scott-Samuel NE (2019) Optimizing colour for camouflage and visibility using deep learning: the effects of the environment and the observer’s visual system. Journal of The Royal Society Interface 16(156)

Johnsen S, Gassmann E, Reynolds RA, Stramski D, Mobley C (2014) The asymmetry of the underwater horizontal light eld and its implications for mirror-based camouflage in silvery pelagic sh. Limnology and Oceanography 59(6):1839–1852

Kondo S, Miura T (2010) Reaction-di usion model as a framework for understanding biological pattern formation. Science 329 5999:1616–1620

Lind O, Henze MJ, Kelber A, Osorio D (2017) Coevolution of coloration and colour vision? Philosophical Transactions of the Royal Society of London Series B, Biological Sciences 372(1724)

Margalef R (1968) Perspectives in Ecology Theory. University of Chicago Press

Marshall J, Johnsen S (2017) Fluorescence as a means of colour signal enhancement. Philosophical Transactions of the Royal Society of London Series B, Biological Sciences 372(1724)

Merilaita S, Scott-Samuel NE, Cuthill IC (2017) How camouflage works. Philosophical Transactions of the Royal Society of London Series B, Biological Sciences 372

Mulero-Pázmány M, Jenni-Eiermann S, Strebel N, Sattler T, Negro JJ, Tablado Z (2017) Unmanned aircraft systems as a new source of disturbance for wildlife: A systematic review. PLoS One 6(6)

Murray JD (1981a) A pre-pattern formation mechanism for animal coat marking. Journal of Theoretical Biology 88(1):161–199

Murray JD (1981b) On pattern formation mechanisms for lepidopteran wing patterns and mammalian coat patterns. Philosophical Transactions of the Royal Society of London Series B, Biological Sciences 295(1078):473–496

Murray JD, Maini PK (1986) A new approach to the generation of pattern and form in embryology. Science Progress 70(280, Part 4):539–553

Nosov KV, Bespalov YG, Vysotska OV, Strashnenko HM, Pecherska AI (2018) Determination of the systemic colorimetric parameters of unmasking rats at videor registration in urban conditions. Bulletin of NTU KhPI Series: New solutions in modern technologies 26 (1302)(2):22–30

Nothdurft HC (2018) Fast identi cation of salient objects depends on cue location. Tech. rep., Visual Perception Laboratory (VPL), Göttingen, Germany

San-Jose LM, Roulin A (2017) Genomics of coloration in natural animal populations. Philosophical Transactions of the Royal Society of London Series B, Biological Sciences 372(1724)

Shannon CE (1948) A Mathematical Theory of Communication. Bell System Technical Journal 27(3):379–423

Shawkey MD, D’Alba L (2017) Interactions between colour-producing mechanisms and their effects on the integumentary colour palette. Philosophical Transactions of the Royal Society of London Series B, Biological Sciences 372(1724)

Sheth R, Marcon L, Bastida MF, Junco M, Quintana L, Dahn RD, Kmita M, Sharpe J, Ros MA (2012) Hox genes regulate digit patterning by controlling the wavelength of a turing-type mechanism. Science 338 6113:1476–1480

Turing AM (1952) The Chemical Basis of Morphogenesis. Philosophical Transactions of the Royal Society of London Series B, Biological Sciences 237(641):37–72

Vysotska O, Balym Y, Georgiyants M, Pecherska A, Nosov K, Bespalov Y (2017) Modeling of a procedure for unmasking the foxes during activities on the elimination of biosafety threats related to rabies. Eastern European Journal of Enterprise Technologies 5(10 (89)):46–54

Zholtkevych GN, Bespalov YG, Nosov KV, Abhishek M (2013) Discrete Modeling of Dynamics of Zooplankton Community at the different Stages of an Antropogeneous Eutrophication. Acta Biotheoretica 61(4):449–465

